# The interaction between dynamic ligand signaling and epigenetics in Notch-induced cancer metastasis

**DOI:** 10.1101/2025.05.19.654987

**Authors:** Tianchi Chen, M. Ali Al-Radhawi, Herbert Levine, Eduardo D. Sontag

## Abstract

Metastatic melanoma presents a formidable challenge in oncology due to its high invasiveness and resistance to current treatments. Central to its ability to metastasize is the Notch signaling pathway, which, when activated through direct cell-cell interactions, propels cells into a metastatic state through mechanisms akin to the epithelial-mesenchymal transition (EMT). While the upregulation of miR-222 has been identified as a critical step in this metastatic progression, the mechanism through which this upregulation persists in the absence of active Notch signaling remains unclear. Here we introduce a dynamical system model that integrates miR-222 gene regulation with histone feedback mechanisms. Through computational analysis, we delineate the non-linear decision boundaries that govern melanoma cell fate transitions, taking into account the dynamics of Notch signaling and the role of epigenetic modifications. Our approach highlights the critical interplay between Notch signaling pathways and epigenetic regulation in dictating the fate of melanoma cells.

## 1 Introduction

Cancer metastasis represents a primary cause of mortality, with the epithelial-mesenchymal transition (EMT) playing a key role in conferring metastatic capabilities upon cancer cells [1–3]. The EMT, characterized by its reversibility, is modulated by a diverse array of environmental cues, EMT-inducing transcription factors (EMT-TFs), and epigenetic regulators [4, 5].

This research specifically targets melanoma, notable for its high resistance to treatment and propensity for metastasis.

The Notch signaling pathway is a critical player in development and disease, including metastasis [6–8]. It regulates cellular differentiation, proliferation, and fate determination [9]. In melanoma, Notch activation, for instance by keratinocytes expressing Notch ligands, can promote metastasis, partly through inhibition of the lineage survival oncogene MITF [10]. Conventionally, Notch activation requires direct cell-to-cell contact. However, melanoma cells can maintain a metastatic phenotype even after losing contact with ligand-expressing cells, suggesting a mechanism for persistence or memory [10].

A powerful approach to describe and resolve the complex interactions and feedback loops involved in genetic and epigenetic regulation is dynamical system modeling [11–13]. In particular, such methods can be applied to regulatory mechanisms involving histone modifications such as H3K4me3 (activating) and H3K27me3 (repressive), which play an important role in EMT and cancer progression [14–16]. We propose that an epigenetic switching mechanism, involving feedback regulation of histone modifying enzymes, underlies the persistence of the Notch-induced metastatic state in melanoma. Specifically, we model the regulation of miR222, whose expression is linked to melanoma metastasis [17] and influenced by Notch [10]. Our model incorporates the competition between NICD (the activated Notch intracellular domain) and MITF for the transcription factor RBPJ, and links this competition to the recruitment of the H3K4me3 demethylase KDM5A [18], thereby influencing the histone state at the miR222 locus. We hypothesize that positive feedback loops in the histone modification system create bistability, allowing the miR-222 locus to be switched to, and maintained in, an active state by a transient Notch signal.

Furthermore, different Notch ligands, such as DLL1 and DLL4, can elicit distinct temporal dynamics of NICD activation – often pulsatile for DLL1 and sustained for DLL4 (Figure 1) [19, 20]. These distinct dynamics can lead to differential activation of target genes [19]. A key question is how these dynamics are interpreted by downstream regulatory circuits. Our model investigates how the proposed epigenetic switch responds to both sustained (DLL4-like) and pulsatile (DLL1-like) NICD inputs, exploring whether the switch exhibits frequency-dependent filtering properties. We find that the epigenetic switch acts as a low-pass filter, which may place high-frequency DLL1 signals at a relative disadvantage compared to sustained DLL4 signals for initiating this specific epigenetic transition.

**Figure 1:**
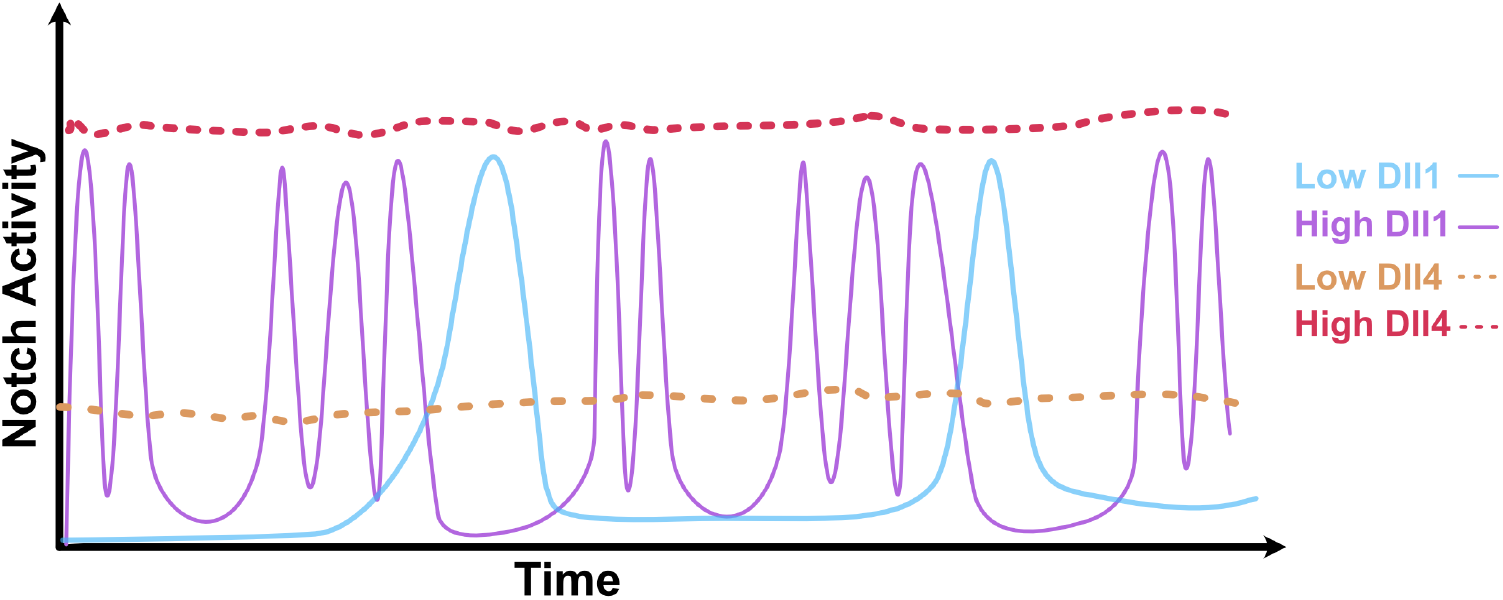
A conceptual view of distinct Notch signaling dynamics induced by DLL1 and DLL4 ligands. Experimental evidence suggests DLL1 ligands can induce pulsatile NICD signals, In contrast, DLL4 ligands tend to induce more sustained signals with an amplitude that increases with the concentration level of the ligand [19, 20]. The characteristics of these signals (amplitude, frequency, duration) can influence downstream cellular responses.

The manuscript is organized as follows: We introduce the computational model linking Notch input competition to miR-222 epigenetic regulation. We then analyze the model’s bistable behavior and its response to sustained and pulsatile NICD signals, characterizing the switching boundaries (amplitude-frequency or *A*-*ω* relationships, where *A* represents signal amplitude and *ω* represents signal frequency) and switching times (*ST* -*ω* relationships, where *ST* represents the time required for state transition). We investigate how altering epigenetic parameters (specifically PRC2 feedback strength) modifies these response characteristics. Finally, we discuss the implications of the model regarding dynamic signal processing, epigenetic memory, and potential therapeutic relevance.

## 2 Methods

### 2.1 Model description

Our model (Figure 2) integrates Notch signaling input with epigenetic regulation of miR-222. It consists of two coupled modules.

**Figure 2:**
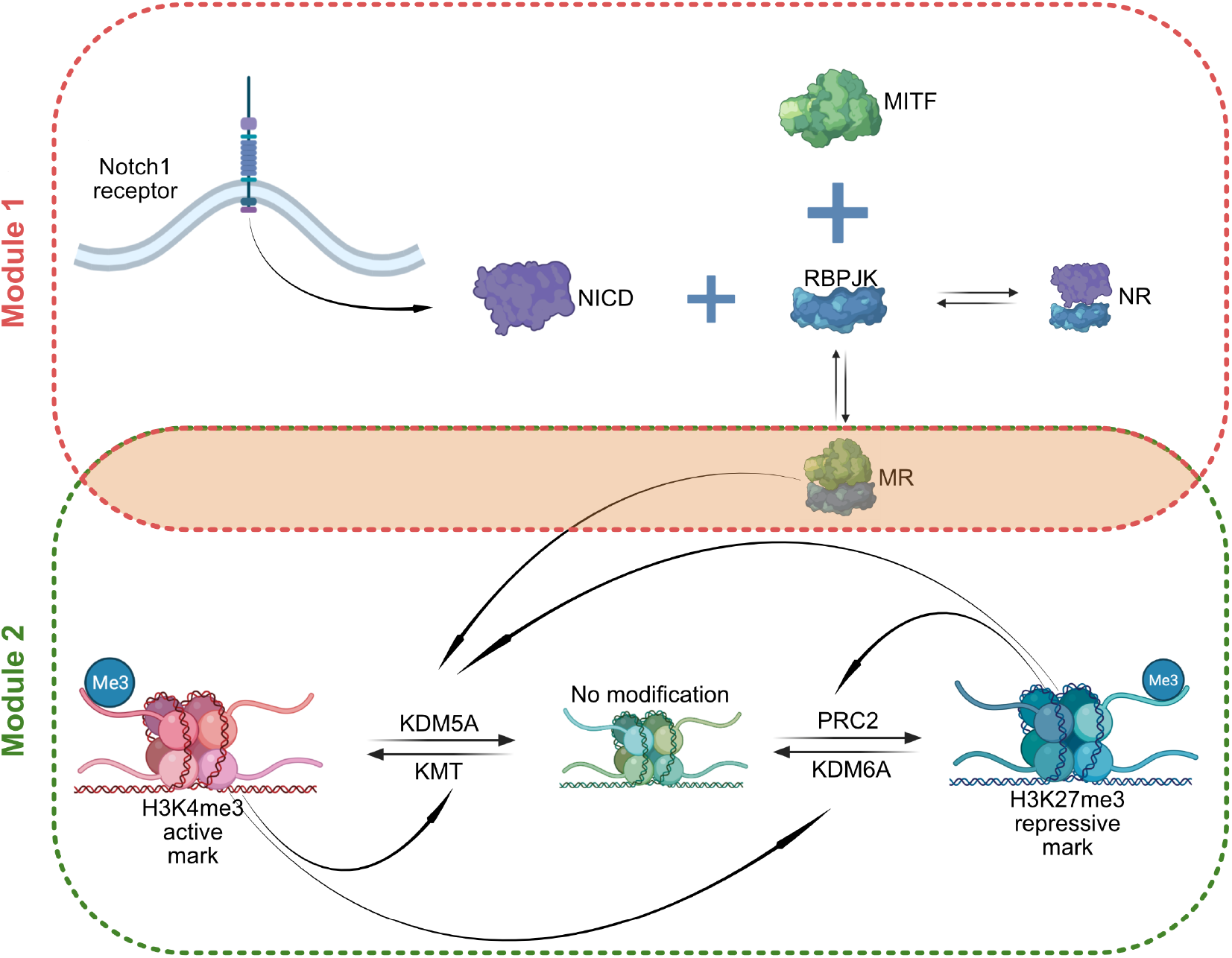
Schematic description of the Notch-EMT model with epigenetic regulations. The model comprises two modules. Module 1 (red dashed region): NICD (N) release is triggered by Notch activation. N competes with MITF (M) for binding to RBPJ (R). High N leads to formation of NR and reduces the level of the MR complex. Module 2 (green dashed region): Represents the epigenetic state of the miR-222 locus via three histone states: H4 (Active, H3K4me3), H27 (Repressive, H3K27me3), and H0 (Unmarked/Void). Transitions are mediated by histone modifying enzymes (KMT, KDM6A, PRC2, KDM5A). Positive feedback (H4 promotes KMT/KDM6A; H27 promotes PRC2/KDM5A) allows for bistability. Coupling: The MR complex enhances KDM5A activity, linking Notch signaling (via MR levels) to the epigenetic state.

#### Module 1: NICD-MITF Competition

Upon Notch receptor activation by a ligand, the Notch Intracellular Domain (NICD, denoted N) is released and translocates to the nucleus. There, it competes with the transcription factor MITF (M) for binding to the DNA-binding protein RBPJ (R). In the absence of NICD, MITF binds RBPJ to form the MR complex. When NICD is present, it binds RBPJ to form the NR complex, thereby reducing the amount of available R and consequently reducing the concentration of the MR complex. The production rate of N, denoted ‘Signal(*t*)’, represents the strength and dynamics of the external Notch ligand stimulus.

#### Module 2: Epigenetic Regulation of miR-222

We model the histone state associated with the miR-222 gene locus using three states: H4 (representing an active state, high H3K4me3), H27 (representing a repressed state, high H3K27me3), and H0. The H0 state represents an unmarked or intermediate chromatin configuration. In our model, this state is assumed to correspond to a basal or low level of miR-222 transcription, distinct from the actively repressed H27 state (low/off miR-222) and the highly active H4 state (high miR-222). It primarily serves as a transient state through which the locus passes during switching between the H4 and H27 states. The transitions between these states are governed by the activity of four types of histone modifying enzymes: KMTs (adding H3K4me3), KDM6A (removing H3K27me3), PRC2 (adding H3K27me3), and KDM5A (removing H3K4me3). Crucially, the model includes positive feedback loops: the H4 state promotes the production/activity of KMT and KDM6A, while the H27 state promotes the production/activity of PRC2 and KDM5A. This double-positive feedback structure can generate bistability between the H4-high and H27-high states.

#### Module Coupling

The two modules are linked via the MR complex. Based on experimental findings [10,18], we assume that the MR complex enhances the production or recruitment of the H3K4me3 demethylase KDM5A to the miR-222 locus. Therefore, high MITF activity (high MR, low NICD) promotes the H27 state (miR-222 repression), while high Notch activity (low MR, high NICD) disinhibits KDM5A recruitment, allowing the feedback loops to potentially switch the system to the H4 state (miR-222 activation).

#### Mathematical model

We formulated the model using chemical reaction networks (CRNs) [21],assuming mass-action kinetics (Table 1).

**Table 1:**
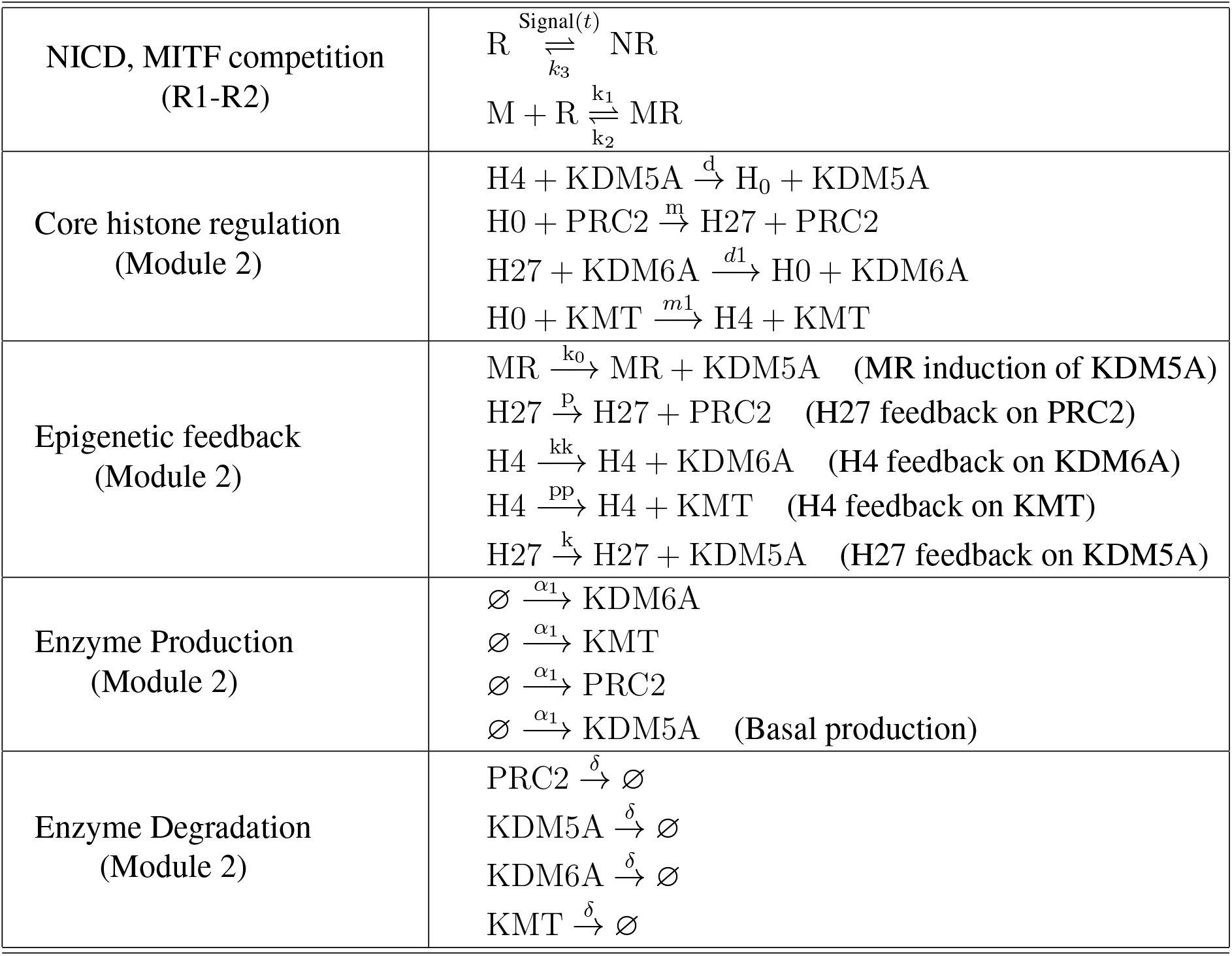
CRN model of the Notch-miR222 circuit. Module 1 is driven by input NICD (N) concentration (Signal(*t*)) and reflects the NICD competition with MITF (M) for RBPJ (R). Module 2 describes the epigenetic regulation of histone states (H4, H0, H27) via enzymes (KDM5A, KDM6A, PRC2, KMT), including feedback loops and basal production/degradation. The modules are coupled via MR-enhanced KDM5A production/recruitment (rate *k*_0_).

**State variables** are the concentrations of the molecules listed in Table 2.

**Table 2:** State Variables of the CRN Model.

| State Variable | Description |
| --- | --- |
| R | Free RBPJ concentration. |
| NR | RBPJ bound to NICD complex concentration. |
| M | Free MITF concentration. |
| MR | MITF-RBPJ complex concentration. |
| H4 | Concentration of loci in the 'Active' histone state (high H3K4me3). |
| H0 | Concentration of loci in the 'Unmarked/Void' histone state. |
| H27 | Concentration of loci in the 'Repressive' histone state (high H3K27me3). |
| KDM5A | H3K4me3 demethylase concentration. |
| KDM6A | H3K27me3 demethylase concentration. |
| PRC2 | H3K27me3 methyltransferase concentration. |
| KMT | H3K4me3 methyltransferase concentration. |

**Parameters** used in the simulations are listed in Table 3, unless otherwise specified. The input signal ‘Signal(*t*)’ directly represents the NICD level, capturing ligand dynamics. For sustained (DLL4-like) input, Signal(*t*) = *A* (constant) during stimulation. For pulsatile (DLL1-like) input, we use a square wave Signal(*t*) = *A*× [1 + sign(cos(*ωt* + *ϕ*))]*/*2 during stimulation, oscillating between 0 and amplitude *A*, with frequency *ω* and initial phase *ϕ*.

**Table 3:**
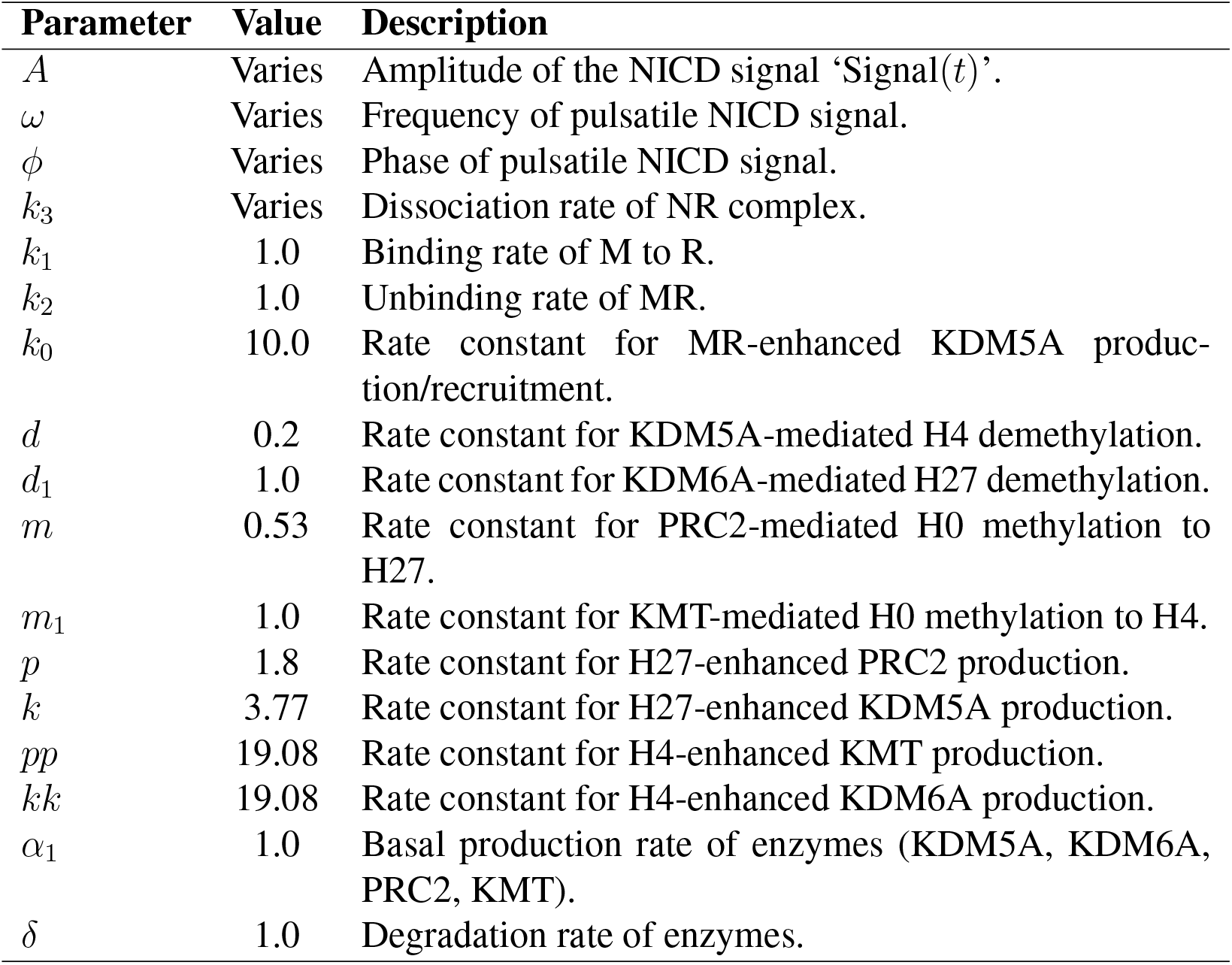
Parameters of the CRN model and default values used. Total amounts of M, R, and Histone sites (H0+H4+H27) are conserved or implicitly set.

| Parameter | Value | Description |
| --- | --- | --- |
| $A$ | Varies | Amplitude of the NICD signal 'Signal($t$)'. |
| $\omega$ | Varies | Frequency of pulsatile NICD signal. |
| $\phi$ | Varies | Phase of pulsatile NICD signal. |
| $k_3$ | Varies | Dissociation rate of NR complex. |
| $k_1$ | 1.0 | Binding rate of M to R. |
| $k_2$ | 1.0 | Unbinding rate of MR. |
| $k_0$ | 10.0 | Rate constant for MR-enhanced KDM5A production/recruitment. |
| $d$ | 0.2 | Rate constant for KDM5A-mediated H4 demethylation. |
| $d_1$ | 1.0 | Rate constant for KDM6A-mediated H27 demethylation. |
| $m$ | 0.53 | Rate constant for PRC2-mediated H0 methylation to H27. |
| $m_1$ | 1.0 | Rate constant for KMT-mediated H0 methylation to H4. |
| $p$ | 1.8 | Rate constant for H27-enhanced PRC2 production. |
| $k$ | 3.77 | Rate constant for H27-enhanced KDM5A production. |
| $pp$ | 19.08 | Rate constant for H4-enhanced KMT production. |
| $kk$ | 19.08 | Rate constant for H4-enhanced KDM6A production. |
| $\alpha_1$ | 1.0 | Basal production rate of enzymes (KDM5A, KDM6A, PRC2, KMT). |
| $\delta$ | 1.0 | Degradation rate of enzymes. |

The corresponding **ordinary differential equations (ODEs)** derived from Table 1 assuming mass-action kinetics are:

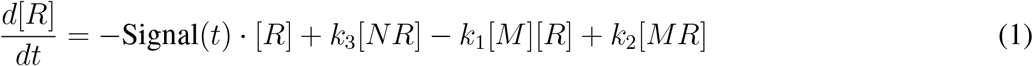

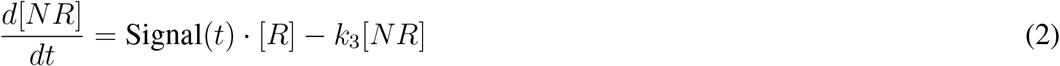

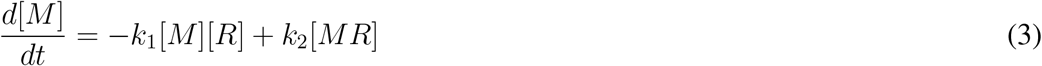

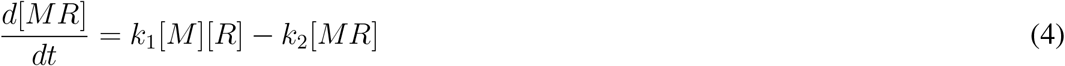

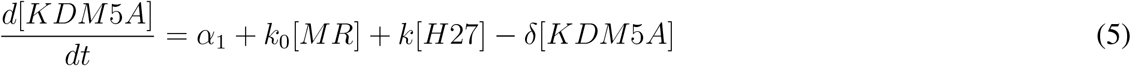

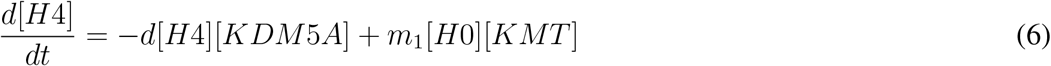

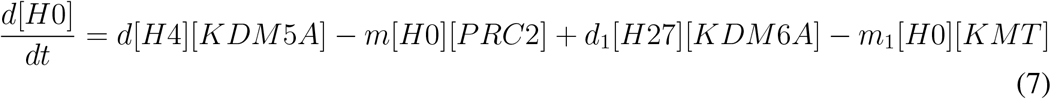

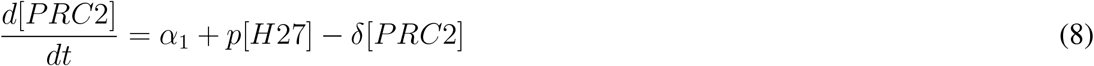

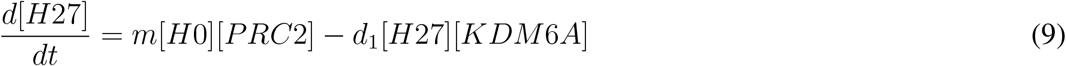

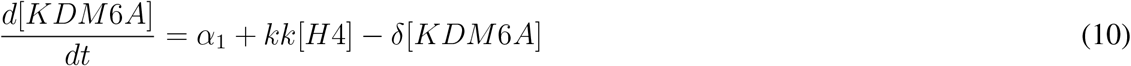

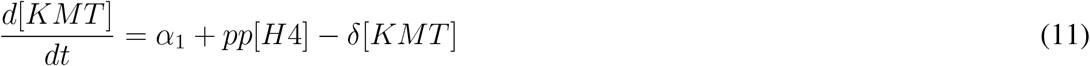

Conservation laws hold for total RBPJ (*R*_*tot*_ = [*R*] + [*NR*] + [*MR*]), total M (*M*_*tot*_ = [*M*] + [*MR*]), and all possible Histone modifications (*H*_*tot*_ = [*H*0] + [*H*4] + [*H*27]).

#### Region of bistability

A key goal of this model is to explain the observed persistence of the metastatic phenotype even after the initiating Notch signal is removed [10]. We hypothesize that this persistence arises from epigenetic memory, mechanistically represented in the model by bistability within the epigenetic module (Module 2). This bistability allows the system to switch between two stable steady states – one corresponding to low miR-222 expression (i.e. H27-high) and another to high miR-222 expression (H4-high) – and remain in the new state after a transient input.

The existence of bistability is fundamental to the model’s ability to exhibit epigenetic memory. We employed standard numerical methods for finding steady states and performing linear stability analysis (based on the eigenvalues of the Jacobian matrix) to identify parameter regimes, including the default set in Table 3, that support this behavior. For these parameters and in the absence of Notch input (N=0), the analysis confirmed that the system exhibits exactly two stable steady states: one corresponding to the repressed state (high H27, low H4) and one corresponding to the active state (high H4, low H27). An unstable steady state typically exists between them. Other potential configurations, such as states dominated by H0, were found to be unstable within the parameter regime supporting bistability between the primary H4-high and H27-high states. In principle it would be possible to search for parameter sets for which the model exhibits tri-stability, but these systems would not be relevant for the biological phenomena we are attempting to capture.

The bistability itself emerges from the positive feedback loops inherent in the epigenetic regulation: the H4 state promoting enzymes for its own maintenance (KMT, KDM6A) and the H27 state similarly promoting its maintenance factors (PRC2, KDM5A), as detailed in Table 1. The existence and parameter range of this bistability critically depend on the strengths of these feedback loops and the rates of histone modification (*k*_0_, *d, m, p, k, pp, kk*, etc.). Therefore, identifying parameter sets that permit bistability is essential for the model to capture the desired memory behavior. We utilized numerical continuation techniques, specifically Homotopy Continuation [22], to explore the parameter space and identify regimes, such as the default parameters listed in Table 3, that yield the necessary two stable steady states in the absence or presence of low Notch input (‘Signal(*t*)’ close to 0).

### 2.2 Algorithms and parameters

The specific parameter values used for simulations are listed in Table 3, unless otherwise stated. These values were selected, using the numerical methods described under “Region of bistability”, primarily to ensure the model exhibits bistability within the epigenetic module (Module 2). This feature is key for representing epigenetic memory based on the positive feedback structure described. While not directly fitted to quantitative experimental data for this specific miR-222 regulatory system in melanoma, the chosen values represent plausible relative strengths and timescales for feedback-driven epigenetic processes often observed in biological circuits [23–26].

Concentrations and kinetic parameters are given in arbitrary units (a.u.). Also, note that our parameter choices establish relative reaction rates that yield the reported dynamics, such as switching events. Experimentally, these events occur on timescales ranging from hours to days, consistent with typical epigenetic processes [19-22]. This sets an approximate value of our time unit as several hours, but a more direct mapping to real time would require calibration via comparison with a specific experimental dataset.

Preliminary analyses suggest the qualitative phenomena of bistability and low-pass frequency filtering are preserved across a range of values for the key feedback parameters (*p, k, pp, kk*), although the precise switching boundaries (e.g., Figure 4) are naturally sensitive to these values. A comprehensive sensitivity analysis across the full parameter space is left for future work (see Discussion).

The external Notch input is represented by ‘Signal(*t*)’ in the ODEs (Eqs. 1-2), modeling the effective concentration of NICD generated. To simulate different ligand inputs observed experimentally [19, 20], we consider the following two scenarios:

- **Sustained (DLL4-like) input:** We use a constant signal, Signal(*t*) = *A*, during the stimulation period.
- **Pulsatile (DLL1-like) input:** We use a square wave oscillating between 0 and amplitude *A*, represented as Signal(*t*) = *A* × [1 + sign(cos(*ωt* + *ϕ*))]*/*2, during stimulation. Here, *A* is the amplitude, *ω* is the frequency, and *ϕ* is the initial phase. This form captures the essential on/off nature of pulsatile signaling.

#### Numerical integration

Simulations were performed in Julia (v1.8+) using DifferentialEquations.jl. The reaction network—encoded with Catalyst.jl—was integrated by the stiff solver Rosenbrock23 (abs./rel. tolerances 10^−6^). Trajectories were initialised in the H27-high steady state, followed by a stimulus of duration Δ*T* (50–100 a.u.). The total runtime was *t*_max_ = 1.5 Δ*T*.

#### Switching criterion

A switch is said to occur when the active-mark species H4 first exceeds the repressive mark H27. Internally, a helper routine scans the numerical solution and returns the first crossing time, denoted *ST* ; if no crossing occurs, the run is classified as non-switching.

#### Phase–independent boundary construction

The DLL1–like input is modelled as a square wave Signal(*t*) = *A* 1 + signcos(*ωt* + *ϕ*)] */*2. For every non–zero driving frequency *ω* we set the initial phase to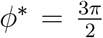, so that Signal(*t*) = 0 for *t <* 0 and the first positive half-cycle starts exactly at the simulation onset *t* = 0. Because any other choice of *ϕ* shifts the waveform leftward in time, *ϕ*^∗^ produces the latest possible arrival of the first activating pulse and therefore represents the mathematically “worst-case” phase for switching.

We then sweep the control parameters over *A* ∈ [0, 300] with unit resolution and *ω* ∈ [0, 2] with step 0.02. For each *ω* we record the smallest amplitude that leads to a switch—this is the conservative threshold *A*^∗^(*ω*). Connecting the points {(*ω, A*^∗^)} yields the phase-independent amplitude–frequency decision boundary shown in Figure 4A. It is important to note that this decision boundary, and the corresponding switching times presented in Figure 4B, were specifically computed for a fixed stimulus duration of Δ*T* = 100 a.u. These boundaries can be dependent on the total stimulus duration; longer durations might allow for switching with weaker or higher-frequency pulsatile signals due to cumulative effects over more cycles, a characteristic not explicitly explored in the current boundary plots. However, further analysis demonstrates that this dependency is weak; for example, increasing the total stimulus duration from Δ*T* = 100 a.u. to Δ*T* = 200 a.u. results in a slight leftward shift of the A-*ω* boundary. Specifically, at a driving frequency of *ω* = 1.0, the minimum amplitude required for switching decreases from approximately *A* = 185 (for Δ*T* = 100 a.u.) to *A* = 180 (for Δ*T* = 200 a.u.). This demonstrates that the system can switch with a slightly weaker signal if the stimulation is applied for a longer total period, confirming the cumulative effect of pulsatile signals over extended durations.

The same simulations provide the switching time *ST* (*A, ω, ϕ*^∗^), defined as the first instant at which the active mark H4 exceeds the repressive mark H27. Grouping these values by amplitude produces the *ST* -*ω* curves in Figure 4B; each curve is an upper envelope valid for all initial phases, because any *ϕ* ≠ *ϕ*^∗^ can only advance the first pulse and shorten the observed switching time.

#### Validation of phase independence

For representative (*A, ω*) pairs we repeated the simulations while sampling *ϕ* uniformly in [0, 2*π*). The maximal switching time and the minimal switching amplitude obtained over the full phase ensemble coincided (within numerical tolerance) with *ST* (*A, ω, ϕ*^∗^) and *A*^∗^(*ω*), respectively, confirming that the reported boundary and *ST* curves are indeed independent of the initial phase.

The GitHub site https://github.com/sontaglab/notch includes the code used to generate the simulations presented in this paper.

## 3 Results

The chemical reaction network (CRN) model presented in Table 1 provides a framework for understanding the dynamics of Notch signaling activation and its influence on the histone state of miR-222. In a previous study [10], researchers observed that melanoma cells could maintain Notch pathway activation and a metastatic phenotype even when not in direct contact with ligand-expressing keratinocytes. This persistence suggests an underlying memory mechanism. The surface of sender cells contains DLL1 and DLL4 ligands, which trigger distinct signaling patterns when Notch is activated: pulsatile signaling is often associated with DLL1, and sustained signaling with DLL4 [19,20]. However, the relationship between the dynamics of Notch signaling in melanoma metastasis and its epigenetic impact on miR-222 remained unexplored.

### 3.1 Notch ligand dynamics determines melanoma cell state transition

In this paper, we focus primarily on the histone state of the miR-222 gene as an indicator of the metastatic melanoma phenotype. We systematically explore the transition between epigenetic states by studying the switching time (ST) of the histone state in the presence of both sustained (DLL4-like) and frequency-modulated pulsatile (DLL1-like) Notch signals. Experimental evidence has shown that induced dynamics of NICD by different Notch ligands can lead to different activation patterns of downstream Notch-targeted genes, which in turn determine cell fate [19]. In the following results, we analyze how ligand dynamics and epigenetic mechanisms coordinate epigenetic state transitions in our model.

#### 3.1.1 DLL4 ligand-induced sustained NICD triggers persistent melanoma metastasis

While the molecular interactions are believed known, the precise dynamical mechanism establishing persistent cellular memory via Notch signaling requires further elucidation. We initially simulated the model using a sustained DLL4-like Notch ligand signal as the external input (Signal(*t*) = *A*). The simulation results qualitatively reproduced the experimental observations of phenotypic persistence [10]. We used a default set of model parameters (given in Table 3) that allowed for bistable histone states and initialized the model from a repressed histone state (H27-high). This setup mimics the experimental finding of high-level repressive histone marks in melanoma in a Notch-free environment [10]. Our model assumes that miR-222 is maintained in a repressed state due to MR-mediated KDM5A activity and the double-positive feedback in the histone methylation circuit. Upon activation of the Notch signaling pathway with sufficient amplitude and duration, the histone state is expected to switch from a repressive state to an activated state.

In Figure 3a, we present an example simulation illustrating how a sustained DLL4-like Notch ligand signal (*A* = 50 applied from *T* = 0 to *T* = 100, so Δ*T* = 100) induces an epigenetic state change for miR-222, potentially leading to an invasive and metastatic state. The histone state of miR-222 is initially repressed (high H27) and then transitions to the active state (high H4) while the signal is active. At *T* = 100, the Notch ligand signal is removed. The simulation demonstrates that the histone state of miR-222 remains in the active state, consistent with experimental observations of persistence. Our simulation results suggest that the model provides a plausible framework at the epigenetic level for explaining the persistence of a high miR-222 state (associated with invasive melanoma cells even after the removal of the Notch signal.

**Figure 3:**
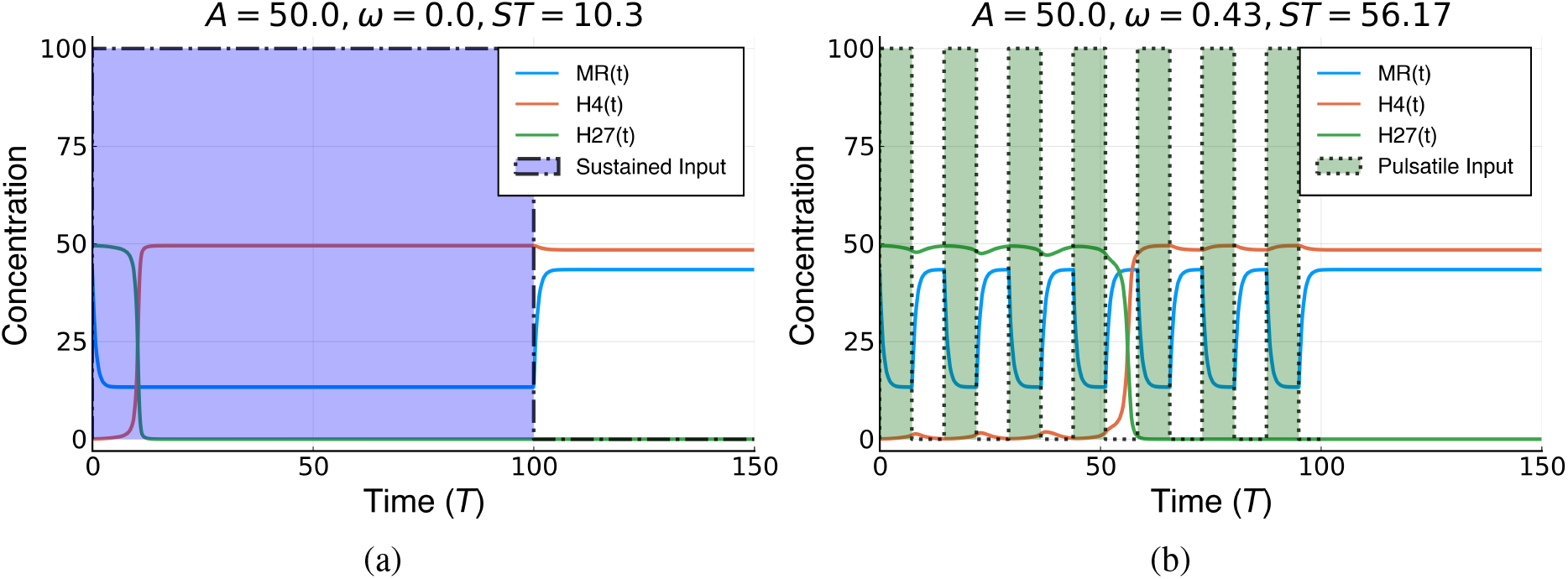
Both sustained (DLL4-like) and pulsatile (DLL1-like) NICD dynamics can induce persistent switching of the miR-222 associated histone state. Simulations start from the H27high state (blue). Signal is applied from T=0 to T=100 (Δ*T* = 100). (a) Sustained input (Signal(*t*) = *A* = 50) triggers a switch to the H4-high state (red), which persists after signal removal. (b) Pulsatile input (Signal(*t*) = 50 × [1 + sign(cos(0.43*t*))]*/*2, *ϕ* = 0) also triggers a persistent switch. Parameters from Table 3.

**Figure 4:**
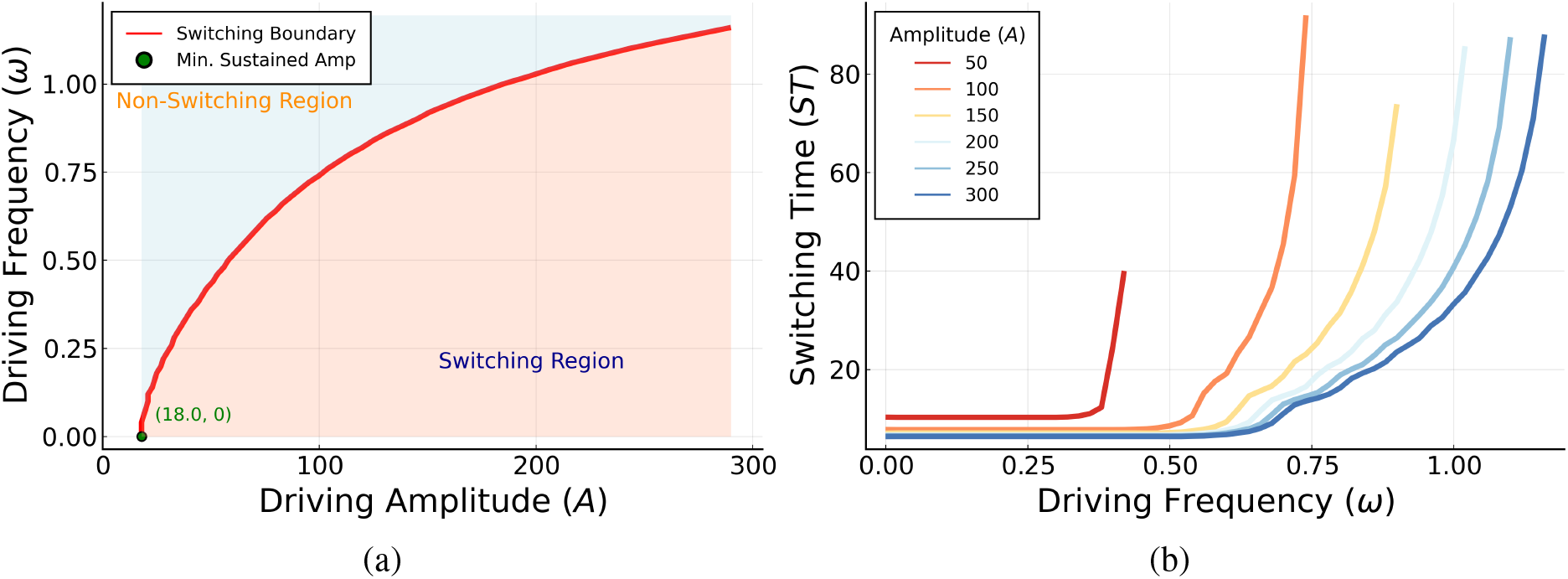
Dependence of epigenetic switching on NICD signal dynamics (phase-independent boundaries computed for duration Δ*T* = 50). (a) Amplitude-Frequency (*A* −*ω*) switching boundary: Minimum amplitude *A* required to ensure switching versus frequency *ω*, regardless of initial phase *ϕ*. Higher frequencies require larger amplitudes (low-pass filter). (b) Switching Time-Frequency (*ST* -*ω*) relation: Maximum switching time *ST* (over all phases *ϕ*) versus frequency *ω* for fixed amplitudes (*A* = 50, 100, 200, 300). *ST* increases sharply above a characteristic frequency for each amplitude. The wiggles along the curves are a numerical artifact.

#### 3.1.2 DLL1 ligand-induced pulsatile NICD can also trigger persistent metastatic melanoma states

The activation of the Notch signaling pathway by DLL1 and DLL4 ligands leads to distinct NICD dynamics (Figure 1). Emerging studies demonstrate the differential effects of DLL1 and DLL4 [19, 27, 28]. This specificity is partly attributed to ligand-receptor interactions modulating these NICD dynamics [19].

So far, we have shown that the DLL4-like ligand can induce a stable change in histone configuration, ultimately leading to the activation of miR-222 (Figure 3a). To understand the influence of DLL1-like ligand input, we modeled pulsatile NICD signals using the square wave form Signal(*t*) = *A* × [1 +sign(cos(*ωt* + *ϕ*))]*/*2 during stimulation. The simulation results, depicted in Figure 3b, reveal that pulsatile induction through DLL1-like signals is also capable of initiating and sustaining miR-222 activation over an extended period, similar to the sustained input from DLL4, given appropriate parameters. The mechanism underlying this response to pulses involves signal integration over time. The epigenetic modification system (Module 2) operates on timescales slower than the NICD fluctuations driven by the pulsatile input. Consequently, while a single short pulse (like the first pulse in Figure 3b, shorter than the switching time seen in Figure 3a) is typically insufficient to cause an irreversible switch, its effect (reducing MR and allowing H4 mark accumulation) partially persists through the ‘off’ phase. Subsequent pulses build upon this lingering effect. If the pulses are sufficiently frequent and sustained over time, the cumulative impact drives the histone state across the threshold for activation, and thereafter the internal positive feedback loops can maintain the H4-high state after the signal ends.

Thus, our analysis confirms that both sustained and pulsatile NICD dynamics can induce longlasting epigenetic changes, leading to stable high-H4 (active miR-222) melanoma cell states, consistent with experimental observations of persistence [10]. Histone transitions triggered by DLL1-induced pulsatile NICD dynamics ultimately exhibit a persistent final pattern similar to that induced by sustained dynamics. However, as discussed further in the context of frequency-dependence in the next section, the efficiency of this integration process is sensitive to the pulse characteristics (amplitude, frequency, duration). Excessively rapid pulsatile

NICD dynamics can hinder the system’s ability to react within each short signaling window, subsequently stalling the transition; this can be thought of as low-pass filtering.

### 3.2 Decision Boundary of Melanoma Cell State Transition

Our research into the epigenetic regulation by Notch ligands establishes that stable changes in the histone state, driven by either sustained NICD signals from DLL4-like ligands or pulsatile signals from DLL1-like ligands. To understand how signal characteristics influence this switch, we computationally determined transition boundaries (minimum amplitude *A* for switching vs. frequency *ω*) as well as the switching times *ST* as a function of (*A, ω*). This analysis aims to clarify how the intrinsic properties of ligand signals collectively influence the thresholds for cell state transitions and the timescale of commitment.

An important aspect of pulsatile signaling is the initial phase (*ϕ*) of the signal. In the computational model, the initial phase affects only the duration of the first pulse, with subsequent pulses being unaffected. This dependence can influence the minimum number of pulses needed to cause a transition. Biologically, this phase must represent the (in general fluctuating) state of the cell at the onset of signal receipt. This initial alignment is likely to be random from cell to cell. To capture the most robust system behavior, one can analyze phase-independent behavior. A phase-independent amplitude threshold (for the *A*-*ω* curve) represents the minimum amplitude required to guarantee switching regardless of the phase, determined by the ‘worst-case’ phase. Similarly, a phase-independent switching time (for the *ST* -*ω* curve) represents the maximum time required to switch across all possible phases.

#### 3.2.1 Amplitude and frequency effects

Figure 4(a) displays the phase-independent *A*-*ω* switching boundary curve. This curve maps out the minimum NICD signaling amplitude (*A*) required to ensure that a stable histone switch occurs at a specific frequency (*ω*) within a fixed duration (Δ*T* = 50), regardless of the initial signal phase *ϕ*. It reveals a non-linear relationship: at low frequencies, a certain minimum amplitude is needed, while at higher frequencies, a larger amplitude is required to achieve switching. This confirms the low-pass filtering nature of the epigenetic switch – it responds less efficiently to high-frequency inputs. This phase-independent threshold reflects the robust signaling strength required to guarantee the epigenetic transition.

#### 3.2.2 Switching Time Frequency (*ST* -*ω*) Relation

Building upon the *A*-*ω* relationship, Figure 4(b) illustrates the phase-independent *ST* -*ω* relationship, showing the maximum switching time (*ST*_*max*_) observed across all initial phases *ϕ* as a function of frequency (*ω*) for several fixed amplitudes (*A*). This represents the “worst-case duration” required for a histone state to shift. We found that this worst-case duration increases with frequency (*ω*) above a certain threshold, reflecting low-pass filtering. Notably, when the frequency is zero (*ω* = 0), the *ST* aligns with what is expected for a sustained DLL4 signal as phase is clearly irrelevant for a constant signal).

More detailed scrutiny of the *ST* -*ω* data reveals that while pulsatile signals from DLL1-like ligands may induce epigenetic changes, they are never faster (especially considering the worst-case phase) than DLL4-like sustained signals in terms of hastening histone state transition times (compare *ST*_*max*_ at *ω* = 0 versus *ω >* 0 for a given *A*). Moreover, the findings underscored in Figure 4(b) convey that at each fixed amplitude, starting from zero frequency, the maximum switching time remains constant up until a definable frequency threshold. Beyond this juncture, the maximum switching time begins increasing rapidly with frequency and eventually diverging at the transition boundary. In essence, the *A*-*ω* and *ST* -*ω* curves reveal how both sustained and dynamic ligand signals can facilitate these epigenetic transitions, albeit within specific parameter ranges. Empirically, this variation in response likely contributes to observations wherein different dynamic signals initiate disparate sets of downstream target [19].

### 3.3 Cooperative control of cell fate by epigenetics and ligand dynamics

Our in-depth examination of the miR-222 gene model has provided key insights into how dynamic ligand signals interact with epigenetic regulation to drive state transitions potentially relevant to EMT in melanoma. Within this framework, histone methylation mediated by PRC2 serves as a key epigenetic control mechanism stabilizing the repressed state. Understanding how PRC2 kinetics influence cell fate transitions provides deeper insight into the coordinated effects of Notch ligand signaling and epigenetic feedback in melanoma progression [29–32].

#### 3.3.1 Epigenetics and Ligand Dynamics Jointly Steer Cell Fate Determination

To gain deeper insight into how PRC2 rate modulates cell fate decisions, we investigate its role in regulating the threshold at which NICD signaling induces miR-222 activation. The precise timing and strength of Notch ligand signals, coupled with epigenetic repression mechanisms, determine whether a melanoma cell switches its epigenetic state.

The PRC2 complex is a crucial histone methyltransferase that catalyzes the deposition of H3K27me3 [33]. Thus, changes in PRC2 rate via parameter *p*) affect the stability of the repressive histone state and alter the Notch signaling threshold required to switch to an active miR-222 state. By understanding this interaction, we can determine how dynamic ligand signaling and epigenetic feedback mechanisms cooperate to define stable cell states [34–36].

#### 3.3.2 PRC2 Rate as a Determinant of Cellular Decision Boundaries

To further elucidate the role of PRC2 rate in cell fate decisions, we investigate how modulating PRC2 activity shifts the epigenetic decision boundary governing miR-222 activation. As displayed in Figure 5, our simulations examine how PRC2 rate affects the amplitude-frequency (A-*ω*) threshold, revealing its role as an epigenetic tuning parameter that controls the sensitivity of miR-222 to Notch signaling.

**Figure 5:**
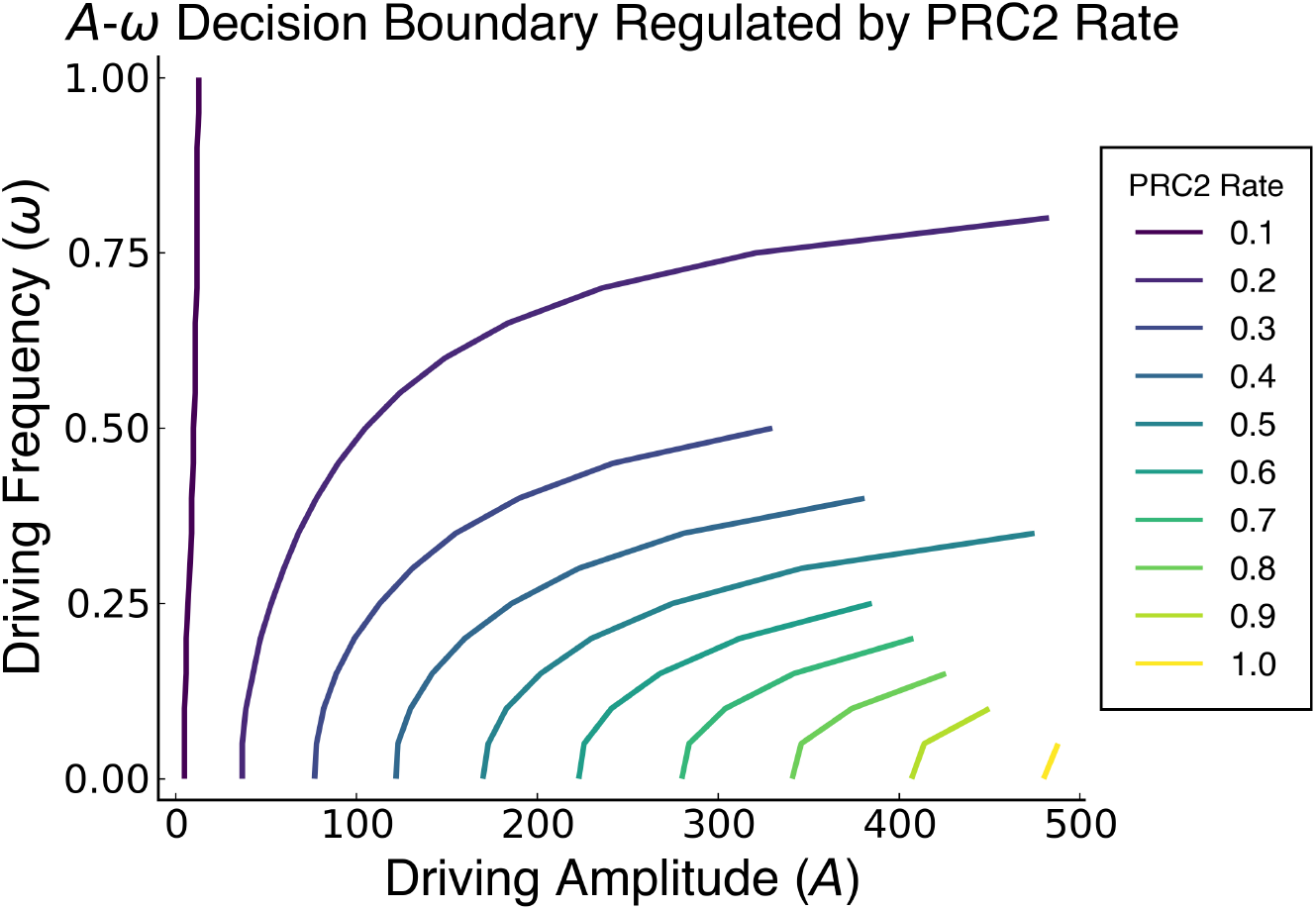
Modulation of the switching boundary by PRC2 feedback strength (*p*). The phase-independent *A*-*ω* boundary (for Δ*T* = 50) shifts downwards and rightwards as *p* increases (from 0.1 to 1.0), indicating that stronger H27-state stabilization requires higher amplitude or lower frequency signals to induce switching regardless of initial phase. Parameter *p* corresponds to PRC2 feedback strength.

By increasing PRC2 rate (parameter *p*), we introduce stronger stabilization of the H27 state, necessitating higher NICD signaling level to overcome the repression and drive gene activation. This effect is particularly evident in the downward and rightward shift of the phase-independent

A-*ω* boundary curve, which indicates that cells with a higher PRC2 rate require a greater Notch signal amplitude or lower frequency to transition to a stable H4-high state. These findings establish PRC2 rate as a critical epigenetic determinant that shapes the Notch-dependent decision boundary, ultimately influencing the stability of epigenetically controlled melanoma cell states.

## 4 Discussion

We have presented a mathematical model integrating Notch signaling input with a bistable epigenetic switch controlling the miR-222 locus. This work was motivated by the observation of persistent metastatic phenotypes in melanoma potentially driven by transient Notch activation [10]. Our model focuses on a core mechanism involving NICD-MITF competition for RBPJ, subsequent regulation of the histone demethylase KDM5A [18], and feedback-driven bistability in H3K4me3/H3K27me3 histone states associated with miR-222.

### Epigenetic Memory and Dynamic Signal Interpretation

A central finding is the model’s capacity for epigenetic memory through bistability. As shown (Figure 3), a sufficiently strong Notch signal, whether sustained or pulsatile, can induce a stable switch to the H4-high (active miR-222) state, which persists after signal withdrawal. This provides a mechanistic explanation for the lasting cellular changes observed following transient Notch signaling [10]. Furthermore, the model reveals how the epigenetic module processes dynamic signals, acting as a low-pass filter (Figure 4). This frequency-dependent response means the system integrates signals over time and is less sensitive to high-frequency pulses compared to sustained or low-frequency inputs. This filtering property offers a way for cells to interpret the distinct temporal dynamics generated by different Notch ligands (e.g., sustained DLL4 vs. pulsatile DLL1 [19, 20]). Such dynamic decoding could underlie the selective activation of downstream targets; for example, genes requiring prolonged signaling to overcome an epigenetic threshold might be preferentially activated by sustained DLL4-like signals, mirroring observations regarding Hes/Hey regulation [19].

### Role of Epigenetic Feedback and Crosstalk

The model’s behavior hinges on the epigenetic feedback loops (Table 1), where histone marks influence the enzymes that modify them [37,38]. The interplay between activating (H3K4me3 via KMT/KDM6A) and repressive (H3K27me3 via PRC2/KDM5A) loops, coupled with the Notch-MR-KDM5A link [18], creates the bistable switch. The significance of epigenetic regulation, including histone methylation and demethylation, in modulating Notch pathway output is well-established [33,39–42]. Our model specifically highlights the role of PRC2, a key regulator of H3K27me3 [43, 44]. Modifying the PRC2 feedback strength alters the signaling threshold required for switching (Figure 5), demonstrating how intrinsic epigenetic factors can tune cellular sensitivity to Notch input. This dynamic interplay between signaling and epigenetics is increasingly recognized as crucial in development and disease, including processes such as EMT [13, 45] and cell fate decisions [46, 47]. While our model focuses on this core axis, Notch signaling operates within a complex network, interacting with pathways involved in metastasis and immunity (e.g., JAK/STAT, PD-1/PDL1) [48–51]. These interactions provide context but are not explicitly included in our current model.

### Therapeutic Implications and Model Generality

The model’s prediction that epigenetic parameters tune sensitivity to Notch signals suggests therapeutic possibilities. EZH2, the catalytic subunit of PRC2, is a target in cancer therapy [52–54], with inhibitors under investigation [34, 55]. Our model implies that modulating EZH2/PRC2 activity could potentially alter the cellular response threshold to oncogenic Notch signaling [16]. The derived non-linear decision boundaries (Figs 4, 5) provide a quantitative framework for exploring such interventions. The underlying motif of a signaling pathway gating a bistable epigenetic switch may represent a general principle [56]. Similar concepts appear in NF-*κ*B-driven epigenetic reprogramming [57] and drug resistance mechanisms [58]. Testable hypotheses arise from the model, such as how manipulating factors affecting ligand dynamics (e.g., glycosylation [59]) or chromatin accessibility (e.g., SWI/SNF activity [60]) might shift the predicted switching boundaries.

### Limitations and Future Directions

While this model provides valuable insights into the interplay between Notch signaling dynamics and epigenetic memory, it represents a simplification of the complex biological reality. The model focuses specifically on the epigenetic state (H3K4me3/H3K27me3 modifications) of the miR-222 locus as a proxy for the metastatic state, omitting the subsequent steps of transcription, translation, and the broader functional consequences characteristic of EMT. Furthermore, the intricate details of Notch signaling, such as cis-inhibition [61] and extensive crosstalk with other pathways relevant to metastasis and immunity, are not explicitly incorporated. The signal dynamics used (sustained constants and idealized square waves) are simplifications of physiological signals. A key strength is that the primary analyses of switching boundaries, including the assessment of PRC2 rate modulation (Figs. 4, 5), considered phase-independent thresholds, reflecting a robust system response.

Future research building upon this work could address these limitations. Incorporating downstream molecular events, such as miR-222 expression levels and their effects, would provide a more complete picture. Exploring the impact of stochasticity in signaling and epigenetic processes, alongside a more thorough analysis of phase averaging for pulsatile inputs, would enhance biological realism. Expanding the model to include interactions with other key signaling pathways involved in melanoma progression is another important avenue. Utilizing more physiologically realistic Notch signal inputs derived from experimental data and performing quantitative comparisons between model predictions and time-resolved measurements would be crucial for validation. Finally, linking the modelś predicted epigenetic switching dynamics with cell-state trajectories observed in single-cell transcriptomic data [62] could offer deeper insights into the temporal regulation of cell fate decisions during Notch-induced metastasis.

## Acknowledgment

HL acknowledges the support of the NSF through the Center for Theoretical Biological Physics, grant no. PHY-2019745 and through DMS-245957. EDS acknowledges the support of AFOSR through grant FA9550-21-1-0289. TC and EDS acknowledge the support of NSF through grant DMS-2052455.

## Data availability statement

All data that support the findings of this study are included within the article.

## Code availability

The simulation code uses Matlab and Python, and all parameters are explained in the paper. All code is available from the following GitHub repository: https://github.com/sontaglab/notch

## Author Contributions

HL and EDS: conceived the study. TC and MAA: performed research and model development. TC: performed the computations. ATC, MAA, HL, and EDS: wrote and edited the paper.

